# Carbapenem-resistant *Klebsiella pneumoniae* lineage CG307 displays urinary tract tropism

**DOI:** 10.64898/2026.04.02.715352

**Authors:** Kyle D. Buchan, Jesus M. Duran Ramirez, Jana Gomez, Timothy R. Cruz, Ender Volkan, Micaela N. Sandoval, Allyson E. Shea, Jennifer N. Walker, Blake M. Hanson

## Abstract

Carbapenem-resistant (CR) *Klebsiella pneumoniae* (*Kp*) are designated by the WHO as a top-priority pathogen due to their antibiotic resistance profiles, capacity to disseminate resistance, and associated mortality. The prototypical CR*Kp* clade, CG258, is associated with acute respiratory infections; however, urinary tract infections (UTIs) caused by CR*Kp* are increasing, and frequently linked to the emergent clade CG307. Notably, CG307 isolates have extensive accessory genomes that may drive adaptation to the urinary tract, including a novel capsule gene cluster and high-affinity urea transporter. In this study, we show that UTIs caused by *Kp* are increasing across the Southern US and that in addition to CG307s circulating within Houston, TX hospital systems, the lineage was also detected in healthcare systems in the broader Gulf Coast region. Characterization of CG307 isolates demonstrate that while these strains exhibit similar mucoviscosity compared to the reference UTI strain TOP52, the lineage displays significantly higher *i)* growth in artificial urine, *ii)* urease activity, and *iii)* UTI in a mouse model. These results suggest that CG307 is spreading across the Southern US and encodes distinct pathogenic features that promote urinary tract tropism, underscoring a need for targeted surveillance and future studies that mechanistically examine the factors that promote UTI.

## INTRODUCTION

Multidrug-resistant (MDR) *Klebsiella pneumoniae* (*Kp*) is one of the primary drivers of the ongoing global antimicrobial resistance (AMR) crisis and is recognized as a major public health threat by the CDC and WHO [1–4]. The WHO’s 2024 Bacterial Priority Pathogens list identifies Carbapenem-resistant (CR) *Kp* as the highest priority pathogen, reflecting its transmissibility, limited preventability, and associated mortality [5]. While CR Enterobacterales (CRE) were rare in the United States in the 20^th^ century, over the last 20 years CREs have increased from ∼1% to >4% of Enterobacterales isolates [6,7]. Furthermore, previous studies indicate that large metropolitan areas within the Northeastern United States report three times as many CRE compared to the Midwest, West, or South, with up to 21% of MDR *Kp* isolates displaying CR in New York City hospital systems [7,8]. More recent reports show CRE is no longer restricted to the Northeast, as the South is reporting rapidly increasing CRE [2,9], with the majority of isolates identified as CR*Kp*. The prevalence of extended-spectrum β-lactamases (ESBLs) such as *bla*_CTX-M-15_ and carbapenemases (including KPC and NDM) among epidemic CR*Kp* lineages worldwide makes treating these infections persistently challenging [10]. The two major clonal groups (CGs) of epidemic CR*Kp* are CG258 and CG307, which consistently harbor a copy of KPC-2 [2,9]. CG258 became established as a primary epidemic strain of CR*Kp* in 2008 and is typically associated with lower respiratory infections (LRIs) [11–13]. However, ours and other’s recent studies demonstrated that the emergent CG307 lineage is increasing in prevalence, associated with high mortality, more frequently isolated from the urinary tract than CG258, and one of the most common lineages circulating in the Southern United States, particularly in Houston, TX [9,14,15]. Notably, CG307 is known to extensively share mobile genetic elements (MGEs) with other clades, further accelerating the spread of AMR to other lineages [9]. These findings are particularly concerning given the global presence of these strains throughout the Americas, Europe, and Asia [16] and the current lack of active surveillance, restricting our ability to track the true prevalence and spread of these MDR lineages.

While *Kp* is typically recognized for its role in causing pneumonia and accounts for a significant proportion of LRIs globally each year [17], the pathogen is more often isolated from the urinary tract due to the sheer number of urinary tract infections (UTIs) that occur annually [18]. Of the estimated 405 million UTIs that occur globally each year [13], 6-12% are caused by *Kp*, which accounts for >28 million cases, making them second only to Uropathogenic *Escherichia coli* (UPEC) [19,20]. Within the United States, a recent report indicated *Kp*-related UTIs sharply increased in from 2011 to 2019, with the highest burden observed in the West South-Central region [15]. Challengingly, of the UTIs caused by *Kp,* >30,000 are caused by MDR strains [21,22]. Notably, our previous study assessing the prevalence of CRE within the United States demonstrated that the MDR CG307 lineage, detected in hospital systems in Houston, TX, were more frequently isolated from the urinary tract compared to the CG258 lineage, which was highly associated with LRIs [9]. Analysis of the accessory genome changes between the CG258 and CG307 lineages identified several acquired genes unique to CG307s [9] including a second capsule synthesis gene cluster (*cp2*), a glycogen synthesis operon (*glgBXCAP*) [23], and a high-affinity urea ABC transporter termed UTS, which consists of UrtA (urea binding protein), UrtB (urea permease), UrtC (urea transporter) and AmiF (formamidase) [9,14]. Together, acquisition of these genes were suggested to impact mucoviscosity, serum resistance (*cp2*), long-term environmental persistence (*glgBXCAP*) [14,24,25], and urinary tract fitness (UTS and *glgBXCAP*) [26–28]; thereby promoting adaptation of CG307 to the urinary tract and contributing to the increased prevalence of *Kp*-UTIs observed in the United States.

In this study, we determined the prevalence of *Kp* UTIs in the Gulf Coast region and assessed phenotypic differences between CG307s, CG258s, and reference UTI strains of *Kp* (TOP52) and UPEC (UTI89) using a range of *in vitro* and *in vivo* assays. Using two health system databases - HealthConnect Texas and USA Health - our data indicate that the prevalence of *Kp* UTIs relative to UPEC is increasing across the Gulf Coast, Alabama, and parts of Mississippi and Florida. Next, we selected 13 representative CG307 isolates for characterization based on the accessory genome, site of isolation, clinical status, and MDR gene carriage. Surprisingly, almost a third of CG307 strains isolated from the urinary tract, including symptomatic UTIs, were observed to have an ∼ 30 kb deletion of the chromosome, which included the type-1 fimbriae operon (*fim*) – a canonical UPEC and *Kp* UTI virulence factor [29–31]. These results suggested that the CG307 lineage may use an uncharacterized pathogenic mechanism for UTI. Thus, growth, mucoviscosity, urease activity, and *in vivo* colonization among these strains were all assessed. While all strains displayed robust growth in rich media, CG307 and CG258 isolates displayed significantly increased growth in artificial urine compared to the reference strain TOP52. Interestingly, mucoviscosity was similar across all strains tested, suggesting the additional capsule synthesis locus (*cp2*) does not impact this trait. In contrast, CG307 strains displayed significantly increased urease activity compared to CG258s and TOP52. Impactfully, CG307 strain C234 exhibited significantly higher bladder colonization compared to TOP52 in a mouse model of UTI. Together, these results suggest that the CG307 lineage is adapted to the urinary tract.

## RESULTS

### UTIs caused by *Kp* are increasing in prevalence across the Gulf Coast region

To investigate the prevalence of *Kp* UTIs across the Gulf Coast region we examined two clinical data sources - HealthConnect Texas and USA Health. HealthConnect Texas is a Health Information Exchange that aggregates electronic health record feeds from over 2,500 facilities across Southeast Texas (hospitals, urgent care centers, outpatient practices, and clinical laboratories) (Fig. 1a). USA Health stands as the only academic health system along the upper Gulf Coast and employs more than 7,200 clinical and nonclinical staff members – including some 180 academic physicians – and serves southern Alabama, southeastern Mississippi, and the western Florida panhandle (Fig. 1a). Impactfully, we observed increasing rates of *Kp* UTIs in both the Houston, TX metropolitan area (HealthConnect Texas) and the Gulf Coast region of Alabama, Mississippi, and Florida (USA Health) between 2017 and 2024 (Fig. 1b). Importantly, the proportion of UPEC – the leading cause of UTIs – remained stable during our observation period, indicating that the observed increase in *Kp* incidence is not due to a change in the rate of UTI. These data are consistent with previous literature reviews and suggest that *Kp* UTIs are continuing to increase in prevalence across the Gulf Coast region [19,32].

**Figure 1.**
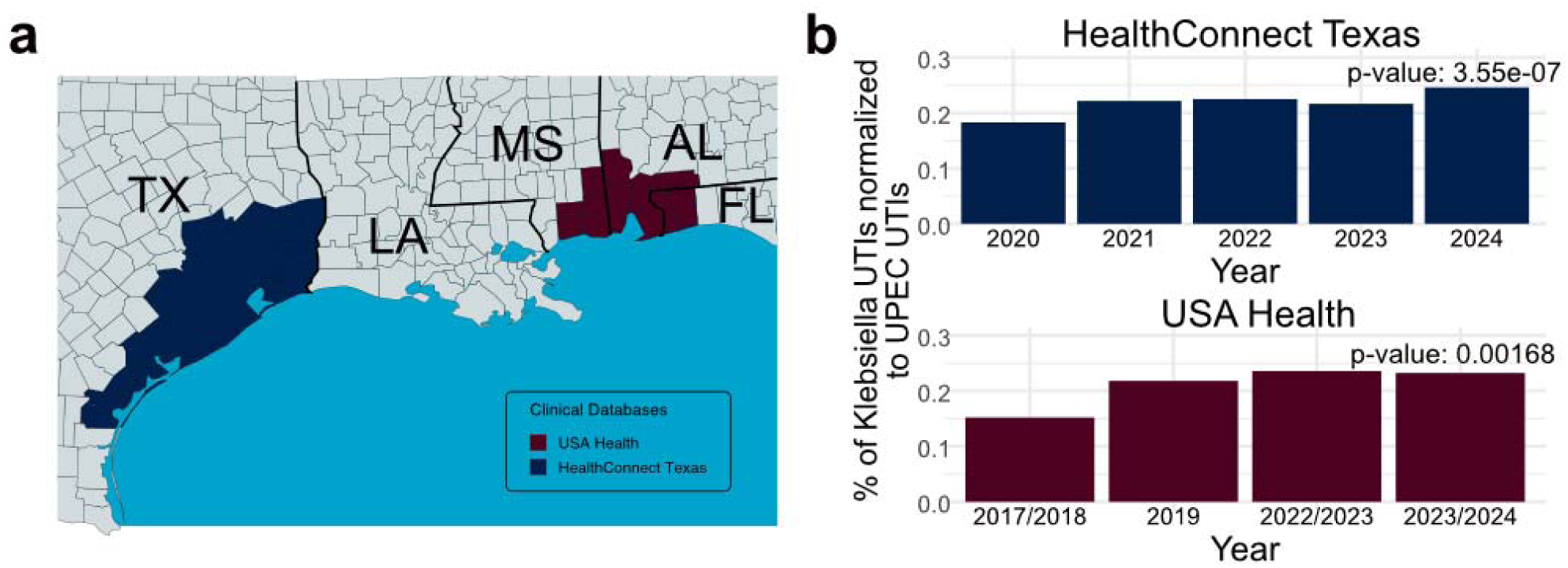
Clinical data from HealthConnect Texas and USA Health showing the *Klebsiella pneumoniae* (*Kp*) urinary tract infection (UTI) incidence over time. **a)** Map depicting the geographic areas from which encounter data were obtained, derived from the HealthConnect Texas (blue) and USA Health (red) databases. **b)** Incidence of *Kp* UTIs compared to uropathogenic *Escherichia coli* (UPEC) UTIs between 2020 and 2024 for HealthConnect Texas and 2017 and 2024 for USA Health. Significance over time was determined using a Chi-square test.

### Genomic characterization of representative CG307 isolates

To characterize the CG307 lineage circulating within the Gulf Coast region between 2017-2024, we selected 13 representative isolates (approximately one third of the total cohort) that were prospectively collected, sequenced, and annotated (CG307 strains listed in **Table 1**) [9]. These CG307 isolates were collected from different anatomical sites (2 respiratory, 11 urinary), represent various clinical statuses (8 colonization, 5 infection), and exhibit variable carriage of carbapenemase genes (3 without, 10 with). A reference-based phylogeny of the 13 selected CG307 strains indicate all isolates are similar and encode a variety of AMR genes, including the KPC-2 carbapenemase (10/13 strains) and at least two copies of the ESBL gene *bla*_CTX-M-15_ (12/13 strains) **(Fig. 2)**. Additionally, as previously reported [9,14] all CG307 strains encoded the UTS and *cp2*, a second, identical K and O loci - the *wzi-173* allele associated with the KL102 locus and the O2v2 locus, respectively **(Fig. 2)**. Notably, ∼38% (5/13) of CG307 isolates exhibited a large scale (30 kb) deletion within the core genome, which was similar to the broader cohort. This deletion included the *fim* operon, which encodes the type 1 fimbrial locus *fim* – an essential *Kp* UTI virulence factor [29] – leaving these strains deficient for type 1 pili (**Fig. 2**). Paradoxically, all strains with this observed deletion either caused symptomatic UTI or were isolated from asymptomatically colonized urinary tracts (**Fig. 2**), indicating that these strains encode alternative mechanisms of urinary tract colonization and/or infection.

**Figure 2.**
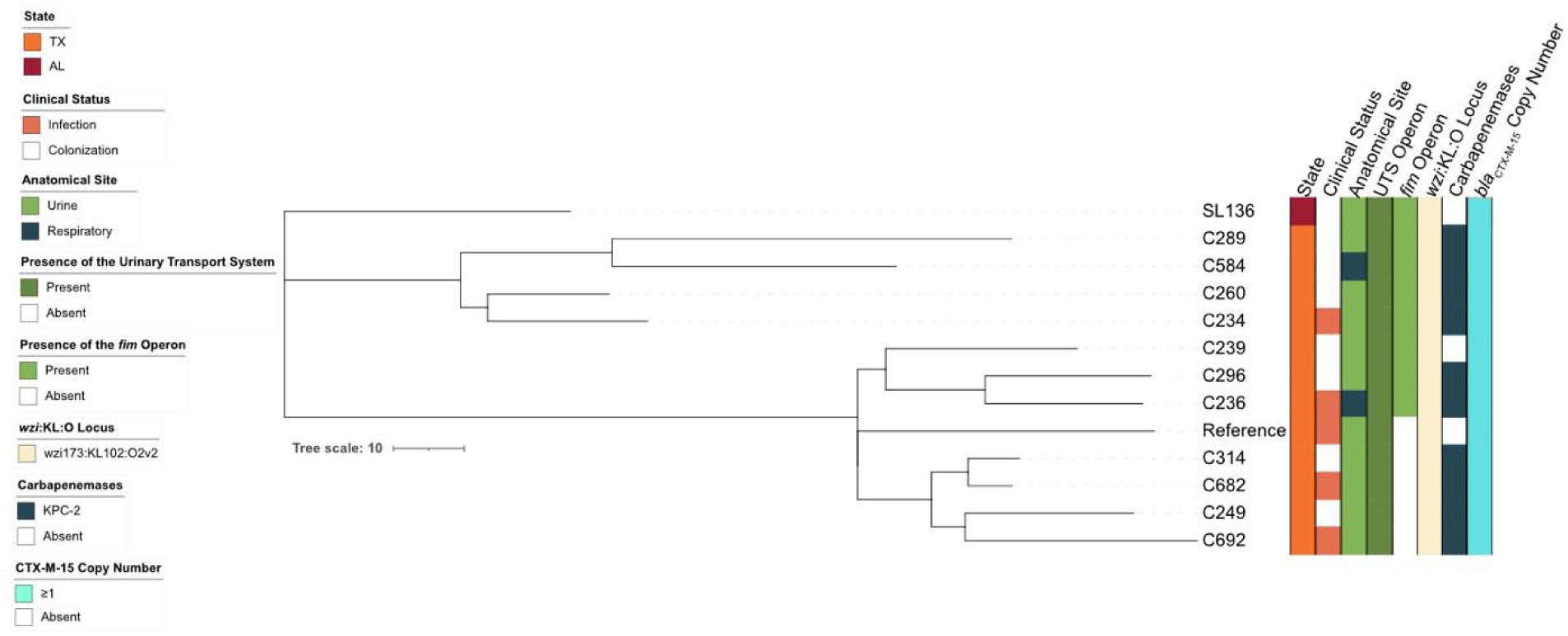
Reference-based phylogenetic tree of CG307 strains used in this study. All strains were collected in Texas (orange) with the exception of SL136 which was from the greater Gulf Coast region (red). Strains were isolated from either symptomatic infections (light orange) or from asymptomatic colonization (white). Strains were cultured from urine (light green) or sputum (dark blue) samples. After sequencing, genomes were queried for the presence of numerous genomic elements. All strains possessed the high-affinity urea transport system (UTS) (dark green). Two thirds of strains had an intact *fim* operon (green), while the remaining third had a complete deletion of the operon (white). All strains possessed an identical *wzi-173* allele and were assigned to the same K and O loci (off-white). Most strains possessed the KPC-2 carbapenemase gene (10/13 – dark blue) while 3 strains did not possess a carbapenemase (white). All strains encoded at least one copy of the extended spectrum β-lactamase (ESBL) gene *bla*_CTX-M-15_ (light blue).

**Table 1.** Strains used in this study. CPE – Carbapenemase-producing *Enterobacteriaecae*.

| Strain ID | BioSample Accession | Genus/Species | Clinical Status | Anatomical Site | Site of Collection | Clonal Group | CPE Status | Carbapenem/ESBL Genes | Reference (PMID) |
| --- | --- | --- | --- | --- | --- | --- | --- | --- | --- |
| C0234 | SAMN15868954 | *K. pneumoniae* | Infection | Urine | Texas | CG307 | CPE | KPC-2, CTX-M-15 | 35357213 |
| C0236 | SAMN15868956 | *K. pneumoniae* | Infection | Respiratory | Texas | CG307 | CPE | KPC-2, CTX-M-15 | 35357213 |
| C0239 | SAMN15868958 | *K. pneumoniae* | Colonization | Urine | Texas | CG307 | Non-CP CRE | CTX-M-15 | 35357213 |
| C0246 | SAMN15868964 | *K. pneumoniae* | Infection | Urine | Texas | CG307 | Non-CP CRE | CTX-M-15 | 35357213 |
| C0249 | SAMN15868967 | *K. pneumoniae* | Colonization | Urine | Texas | CG307 | CPE | KPC-2, CTX-M-15 | 35357213 |
| C0253 | SAMN15868971 | *K. pneumoniae* | Colonization | Urine | Texas | CG258 | CPE | KPC-2, SHV-12 | 35357213 |
| C0260 | SAMN15868976 | *K. pneumoniae* | Colonization | Urine | Texas | CG307 | CPE | KPC-2, CTX-M-15 | 35357213 |
| C0262 | SAMN15868977 | *K. pneumoniae* | Infection | Respiratory | Texas | CG258 | CPE | KPC-2, SHV-12 | 35357213 |
| C0289 | SAMN15868993 | *K. pneumoniae* | Colonization | Urine | Texas | CG307 | CPE | KPC-2, CTX-M-15 | 35357213 |
| C0296 | SAMN15868999 | *K. pneumoniae* | Colonization | Urine | Texas | CG307 | CPE | KPC-2, CTX-M-15 | 35357213 |
| C0314 | SAMN15869015 | *K. pneumoniae* | Colonization | Urine | Texas | CG307 | CPE | KPC-2, CTX-M-15 | 35357213 |
| C0584 | SAMN15869030 | *K. pneumoniae* | Colonization | Respiratory | Texas | CG307 | CPE | KPC-2, CTX-M-15 | 35357213 |
| C0682 | SAMN15869049 | *K. pneumoniae* | Infection | Urine | Texas | CG307 | CPE | KPC-2, CTX-M-15 | 35357213 |
| C0692 | SAMN15869053 | *K. pneumoniae* | Infection | Urine | Texas | CG307 | CPE | KPC-2, CTX-M-15 | 35357213 |
| SL136 | SAMN54466988 | *K. pneumoniae* | Infection | Urinary cathete | Alabama | CG307 | Non-CPE | CTX-M-15 | This Study |
| TOP52 | SAMN02767132 | *K. pneumoniae* | Infection | Urine | Washington | CG152 | Non-CPE | - | 18092884 |
| UTI89 | SAMN00000110 | *E. coli* | Infection | Urine | Michigan | - | - | - | 18411285 |

### CG307 isolates display augmented growth in artificial urine

To determine fitness phenotypes in the urinary tract, CG307 isolates were grown in nutrient-rich (LB) and nutrient limited artificial urine (AU) media. AU media mimics human urine and has been used previously to investigate Gram-positive uropathogens [26]. In addition to our 13 selected CG307 isolates, we included TOP52 (a well-characterized CG152 uropathogenic *Kp* cystitis strain [33], UTI89 (a UPEC cystitis strain [30]), and two CR*Kp* strains from the CG258 lineage (C253 and C262 [2]. All strains exhibited robust growth in LB broth **(Fig. 3a, b)**. Notably, in AU media, 2/13 isolates displayed significantly higher growth compared to TOP52, and 8/13 CG307 strains grew similarly to UTI89. **(Fig. 3c, d)**. These results suggest that MDR *Kp* strains effectively grow within the nutrient-limited urinary tract environment.

**Figure 3.**
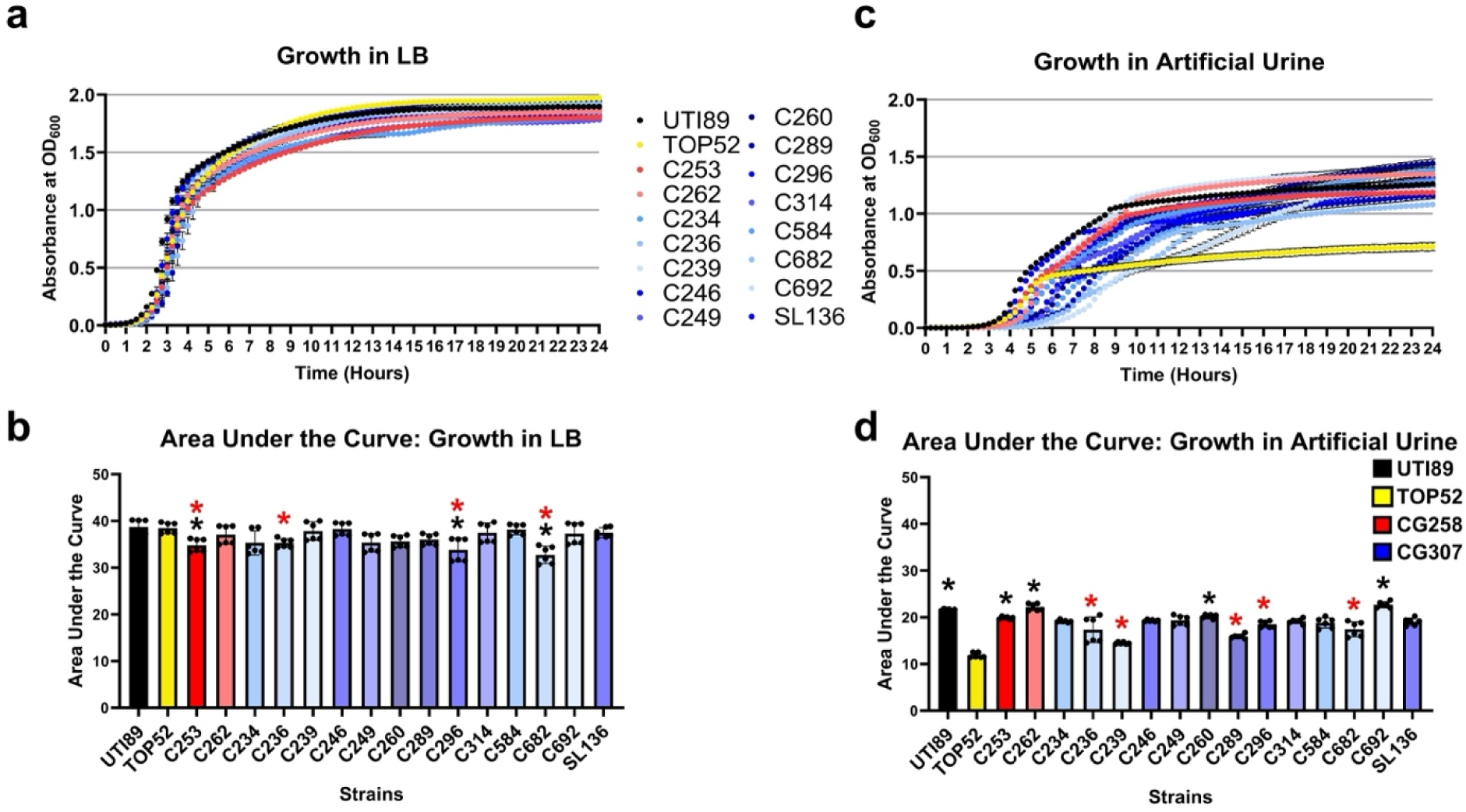
Growth of multidrug resistant (MDR) *Klebsiella pneumoniae* (*Kp*). CG307 and CG258 strains, and control strains TOP52 and UTI89 were grown in **(a, b)** rich media (LB) and **(c, d)** artificial urine media. Significance was determined with a Kruskal-Wallis and Dunn’s test, with black stars indicating comparison against strain TOP52 and red stars indicating comparison against strain UTI89; *p* = *:<0.05. All experiments were performed with two biological and three technical replicates.

### Mucoviscosity is similar across MDR *Kp* strains

The excessive production of capsule by *Kp* - known as hypermucoviscosity – is associated with hypervirulent *Kp* (hv*Kp*) strains [14,34,35]. To determine if the acquisition of a second capsule synthesis gene cluster (*cp2*) by CG307s – which is not present in the CG258 or CG152 lineages – impacts the capsule production of CG307 strains, mucoviscosity was assessed as previously described [36]. TOP52 exhibited an intermediate level of mucoviscosity close to the mean of all tested strains, and 9/13 CG307s displayed equivalent mucoviscosity to TOP52 **(Fig. 4a)**. Four CG307 strains - C239, C314, C682 and C692 - exhibited significantly less mucoviscosity than TOP52. Group-level analyses showed no significant association between mucoviscosity and control (UTI89 and TOP52), CG258, or CG307 groups **(Fig. 4b)**. The similar levels of mucoviscosity displayed by all strains tested suggests that the acquisition of a second, additional capsule synthesis gene cluster (*cp2*) does not significantly impact mucoviscosity within this lineage.

**Figure 4.**
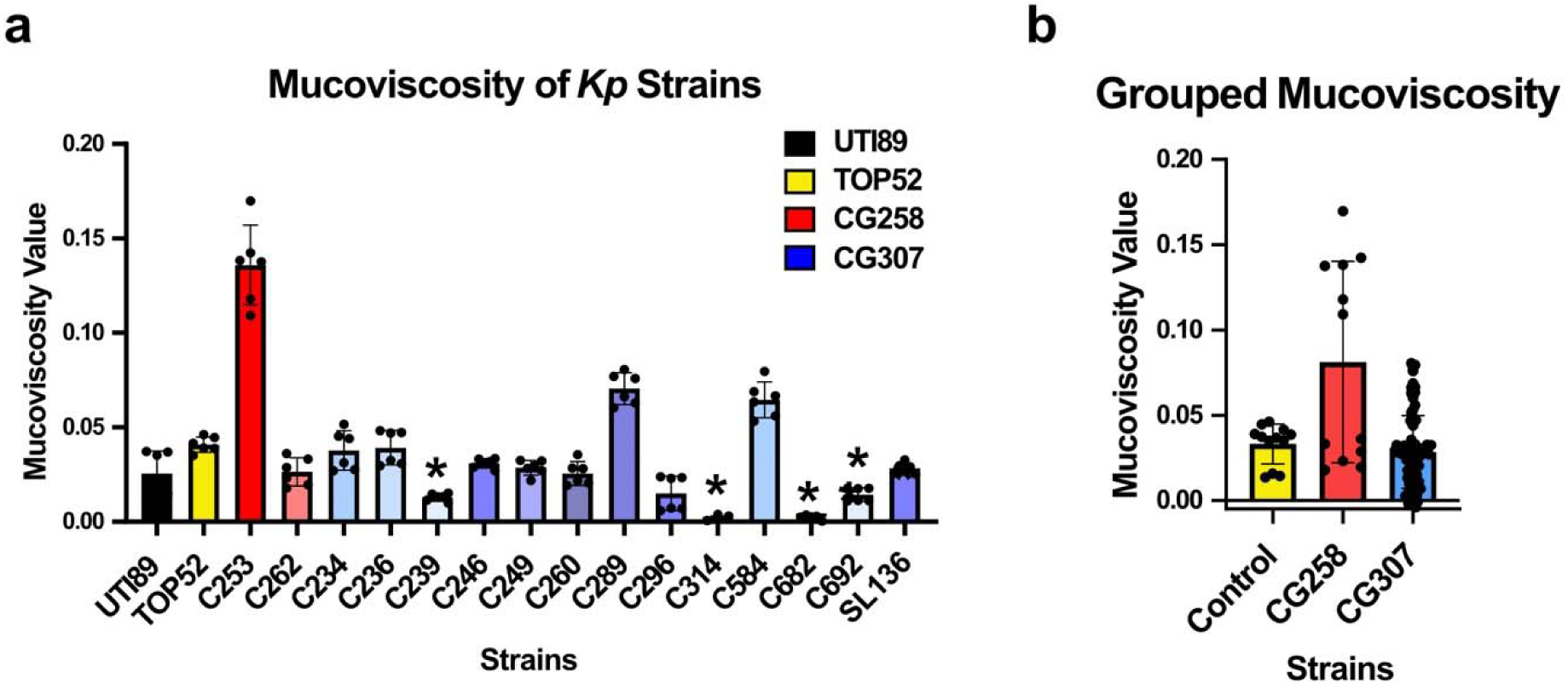
Mucoviscosity of multidrug resistant (MDR) *Klebsiella pneumoniae* (*Kp*). CG307s and CG258s strains, and control strains TOP52 and UTI89 were assessed for mucoviscosity **a)** across individual strains and **b)** grouped. Significance was determined with **a)** a Kruskal-Wallis and Dunn’s test with stars indicating comparison against the TOP52 strain (*p* = *: <0.05) or **b)** a linear mixed effects regression model. Data are reported as the average of three technical and two biological replicates with error bars indicating the standard deviation.

### CG307 isolates display increased urease activity

The acquisition of a high-affinity urea ABC transporter, UTS, suggested urea uptake and hydrolysis may be altered in CG307 [9,14]. Thus, urease activity within all strains was assessed as previously described [37] **(Fig. 5a)**. UPEC strain UTI89 was included as a urease-negative control and TOP52 was included as a comparison for CG307 strains. Among the CG307 isolates, most displayed a similar level of urease activity to TOP52 (9/13). However, almost a third (4/13) of the CG307 isolates displayed significantly higher urease activity compared to TOP52 **(Fig. 5b)**. In grouped analyses, a significant increase in urease activity was observed amongst CG307 strains compared with CG258 strains **(Fig. 5c)**. When CG307 isolates were grouped based on infection site, there was no difference in urease activity in urinary tract isolates (15/17) compared to respiratory isolates (2/17) **(Fig. 5d)**. Together, these data demonstrate that the CG307 lineage (regardless of isolation site) displays higher urease activity compared with the prototypical TOP52 and MDR *Kp* clade CG258.

**Figure 5.**
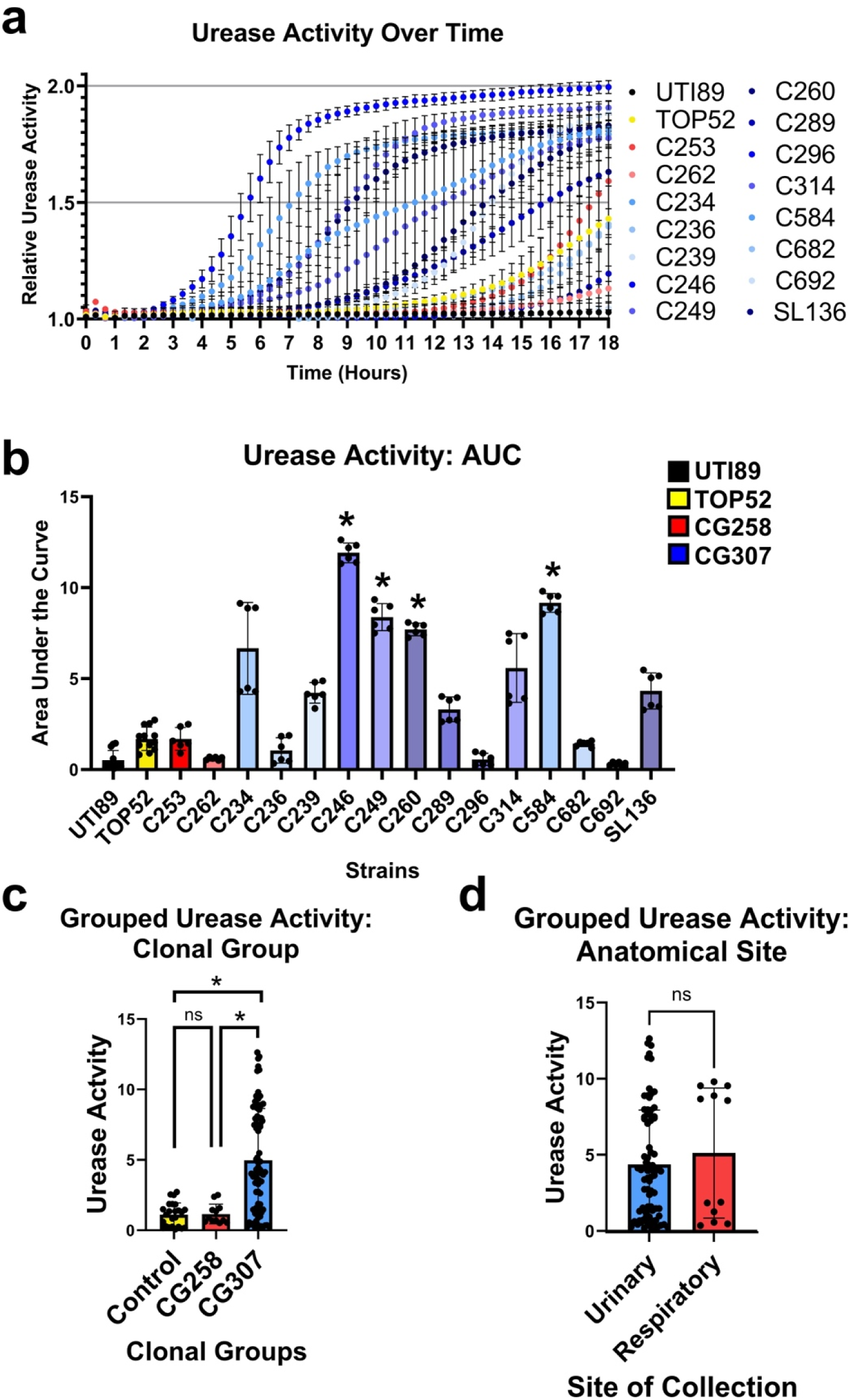
Urease activity among multidrug resistant (MDR) *Klebsiella pneumoniae* (*Kp*) CG307s and CG258s, and the control strains TOP52 and UTI89. **a**) Urease activity of all strains over an 18-hour time course. **b)** Area under the curve of urease activity from all strains over an 18-hour time course. Significance was determined with a Kruskal-Wallis and Dunn’s test, stars show the strain compared against TOP52. **c)** Urease activity by clonal group, groups were compared using a linear mixed effects regression test, *p* = *: <0.05. **d)** Urease activity by site of isolation, compared using a Chi-squared test, *p* = ns: >0.05. Performed with two biological replicates in technical triplicate.

### CG307 strain C234 results in robust UTI

To determine whether CG307 isolates can cause UTI in a preclinical mouse model, we assessed the representative MDR CG307 strain C234. UTI89 and TOP52 were included as positive controls and an isogenic TOP52 mutant with a *fim* operon deletion was included as a negative control [29]. UTI89 robustly colonized the bladder and disseminated to the kidney at 1 day post infection (dpi), as expected (**Fig. 6a, b**). TOP52 displayed significantly lower bladder and kidney colonization compared to UTI89, while the TOP52Δ*fim* mutant was effectively cleared from the bladder and kidneys of mice, as previously reported [29]. Strikingly, MDR CG307 strain C234 displayed significantly higher bladder burden compared with TOP52 and was similar to UTI89 at 1 dpi **(Fig. 6a)**. These findings indicate that MDR CG307 strain C234 robustly colonizes the bladder and outperforms the prototypical *Kp* UTI strain TOP52.

**Figure 6.**
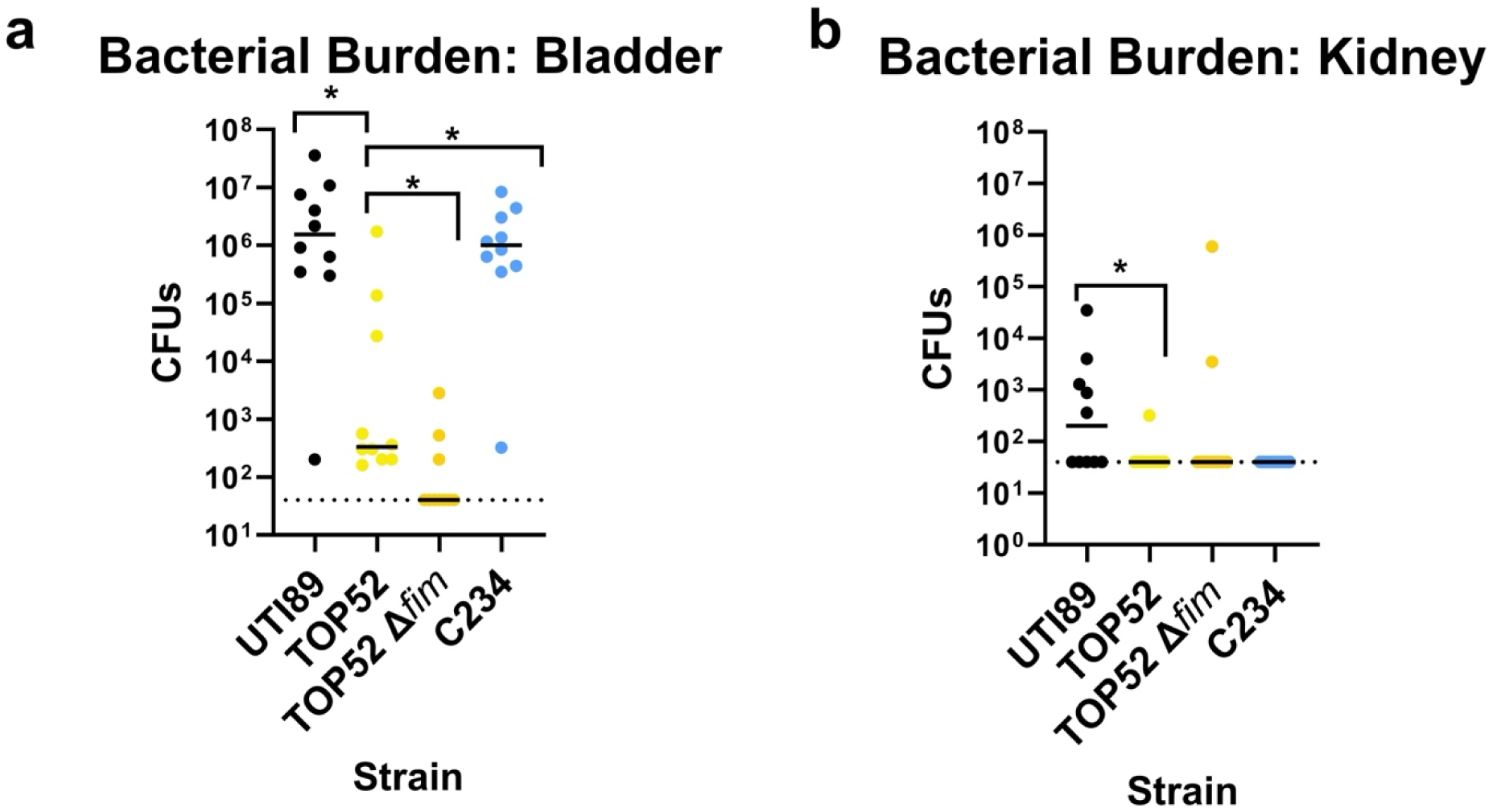
Mouse model of multidrug resistant (MDR) *Klebsiella pneumoniae* (*Kp*) urinary tract infection (UTI) at 1 day post infection (dpi). Cystitis reference strains UTI89 (uropathogenic *Escherichia coli*) and TOP52 (*Kp*), a TOP52 isogenic *fim* operon mutant strain, and an emergent MDR CG307 strain - C234 - were assessed in a mouse model of UTI. Bacterial burden was assessed within **a)** the bladder and **b)** the kidneys at 1 dpi. Dashed lines indicate the limit of detection. Significance was determined with a Mann-Whitney U test, *p* = *: <0.05.

## DISCUSSION

Over the last 20 years, CRE infections have increased in prevalence in the US, with CR*Kp* accounting for a majority of these [2,5,38]. The global spread of two epidemic CR*Kp* lineages, CG258 and CG307, substantially contributed to the increased disease burden in the US [9,14]. Challengingly, CR*Kp* infections are continuing to rise, with the highest increase observed between 2011 and 2019 in the West South-Central census division, which includes Texas [15]. Ours, and others, previous studies investigating the prevalence of MDR *Kp* indicate that the newly emergent CG307 lineage was found to be more frequently isolated from the urinary tract, while the CG258 lineage predominates in the respiratory tract [9,12,13]. Furthermore, CG307 strains causing UTIs are associated with high overall resistance and inappropriate empiric therapy, resulting in increased morbidity and mortality [39–42]. While some studies report an increase in *Kp* UTI and other studies suggest MDR *Kp* UTIs are also increasing, there remains a paucity of data investigating overall *Kp* UTI rates or characterizing MDR *Kp* UTI strains.

In this study, we used two clinical data sources – HealthConnect Texas and USA Health – to determine that the incidence of *Kp* UTI increased between 2020 and 2024 for the Greater Houston area and between 2017 and 2024 for the adjacent region of the Gulf Coast covering Southern Alabama. This trend contrasts with the incidence of UPEC UTI, which remained steady during the same period. Additionally, while we previously reported the presence of the CG307 lineage within the Greater Houston, TX area [9,15], we report here that the lineage was also detected in other regions of the Gulf Coast. While this work provides strong evidence to show that *Kp* UTI incidence is increasing and the CR*Kp* lineage CG307 is continuing to spread across the Gulf Coast region, future studies that prospectively collect and sequence MDR *Kp* UTI isolates are critically needed to determine whether observed increases in *Kp* UTIs correlate with the spread of the CG307 lineage.

Our previous study reported four hierarchical population structures of CG307 strains [9]. However, clade I, which shares a paraphyletic relationship with the globally disseminated clade II, no longer appears to be circulating and was consequently excluded from this study [43]. As observed previously, the remaining three clades consist of one globally disseminated lineage and two Houston-specific, closely related clades, which is accurately reflected in the representative isolates selected characterization in this study. Despite phylogenetic separation, these strains have a highly homogeneous profile of virulence factors and AMR determinants. Impactfully, several acquired genes identified within the CG307 lineage may be associated with urinary tract fitness and/or pathogenesis [9,14]. Specifically, all CG307s encode a second capsule synthesis gene cluster (*cp2*), a glycogen synthesis operon (*glgBXCAP*) [23], a high-affinity urea ABC transporter termed UTS (UrtABC/AmiF) [14], and a fimbrial gene cluster with homology to the pi-family (π-pili) [9,14]. The *cp2* gene cluster may alter mucoviscosity by overproducing capsule and contributing to a hypermucoviscous phenotype, which elsewhere is associated with hypervirulent *Kp* strains [14,35]. Glycogen synthesis is uncharacterized in *Kp*, however in *E. coli* it may aid survival under nutrient-limiting conditions and promote long-term survival outside of the host [14,24]. The high-affinity urea ABC transporter (UTS) is a gene cluster consisting of UrtA (urea binding protein), UrtB (urea permease), UrtC (urea transporter) and AmiF (formamidase), and in tandem with the urease enzyme may confer a selective advantage in the urea-rich environment of the urinary tract [14,44]. The π-pili is an uncharacterized virulence factor with homology to the pap-pilus of UPEC, a prominent virulence factor that enables ascending infection to the kidneys during UTI and may employ a unique pathogenic mechanism during CR*Kp* UTIs [45]. The acquisition of these genes, which are not present in CG258, combined with the strong association of CG307 with the urinary tract, suggests that this lineage has evolved specific tissue tropism.

Strikingly, in-depth genomic analysis of a strain set of 48 CG307 isolates collected previously determined that >20% of isolates lost the *fim* operon - which encodes type-1 fimbriae - as part of a 30 kb chromosomal deletion [9]. Type-1 fimbriae are the canonical virulence factor in UPEC and *Kp* [29–31]. The pilus is critical for mediating initial adhesion to and invasion of bladder epithelial cells, as well as subsequent formation of intracellular bacterial communities during UTI [29–31,46,47]. Notably, the isolates that lost the *fim* operon were overwhelmingly isolated from the urinary tract, with a majority causing symptomatic UTI [9]. This finding suggests that type-1 pili are not essential for CR*Kp* CG307 UTI in some humans and that the lineage has alternative or acquired additional mechanisms that promote urinary tract tropism, such as increased growth, enhanced urea metabolism, or another adhesin. Notably, the CG307 lineage encodes six other pili loci, including π and *mrk* (type 3 fimbriae). Type 3 fimbriae have been shown to be important for catheter-associated UTI (CAUTI) [48]. Yet, few studies have characterized the virulence determinants encoded by CG307 strains, and no study has assessed this lineage for pathogenesis in the urinary tract.

In this study we begin to dissect the fitness and pathogenic mechanisms of the epidemic CG307 lineage in the urinary tract environment. Growth in standard rich media demonstrates that all CG307 strains tested display similar growth patterns to canonical reference UPEC and *Kp* UTI strains. However, growth in artificial urine media demonstrates that the CG307 lineage displays robust growth, which was comparable to the UPEC UTI89 isolates. Notably, many CG307 isolates also displayed significantly higher growth compared to *Kp* TOP52, which exhibited the weakest growth among tested strains. These results suggest that CG307 strains can utilize the limited nutrients, such as urea, within the urinary tract to support robust growth. Future studies will focus on dissecting the fitness advantages provided by the CG307 accessory genome, including the novel synthesis operon (*glgBXCAP*) as well as the high-affinity urea ABC transporter (UTS).

The acquisition of a second capsule locus (*cp2*) within CG307 suggested that these strains may display altered mucoviscosity phenotypes. The excessive production of capsule – termed hypermucoviscosity (HMV) – has been linked to hv*Kp* [25,35]. Notably, the genes *rmpA* and *rmpA2*, which are common to hypervirulent lineages of *Kp sensu stricto*, contribute to HMV phenotypes [49]. HMV is verified with a simple phenotypic assay in which an inoculating loop is lifted from a surface colony producing a viscous ‘string’ >5mm in length [34,35]. However, invasive hv*Kp* strains that do not pass the ‘string test’ have also been reported [35,50,51], suggesting that HMV is not a defining feature of hypervirulence. Our prior study noted that the CG307 lineage does not possess the HMV genes, *rmpA* and *rmpA2* [9]. Furthermore, our mucoviscosity results here suggest that there is no correlation between clonal group and mucoviscosity among the isolates tested. Additionally, we observed a significant decrease in mucoviscosity in several CG307 strains compared with TOP52. Together, these results suggest that mucoviscosity is not a defining trait of this lineage despite possessing two distinct capsule synthesis gene clusters. Future work investigating the role of the two capsule synthesis gene clusters *cps* and *cp2* on mucoviscosity and/or UTI are needed to understand how this duplication may contribute to disease.

The near-universal carriage of the urease operon within *Kp* along with the acquisition of a novel UTS within the GC307 lineage led us to investigate urease activity [9,14]. Urease hydrolyzes urea into ammonia and carbon dioxide [28], and is a rich source of nitrogen within the urinary tract. The breakdown of urea into ammonia releases stored nitrogen, which can be used in metabolism to support microbial growth [44]. Notably, urease only releases stored nitrogen, it does not allow strains to scavenge environmental nitrogen released from the ammonia by extracellular urease. Nitrogen scavenging depends on other transport systems [52]. Interestingly, the acquisition of the UTS within CG307s suggests that this lineage may more efficiently take up urea, which can then be hydrolyzed by intracellular urease, thereby releasing more nitrogen for growth. However, studies in other urease-producing uropathogens, including *Proteus mirabilis*, *Morganella morganii*, and *Staphylococcus saprophyticus* indicate the enzyme is significantly more impactful for its role in virulence [44]. Specifically, the release of ammonia by urease alkalinizes the urinary tract environment and promotes formation of struvite and carbonate apatite crystals [44]. These crystals become embedded in biofilms, promote persistence, increase tissue damage [27,28], and are particularly troublesome in the context of CAUTIs where crystalline precipitates can cause catheter blockages and subsequently device failure [28,53,54]. Despite the critical role of urease in pathogenesis of other urease-producing uropathogens, the enzyme is virtually unstudied in *Kp* UTI. Our results reported here demonstrate that the CG307 lineage displays significantly higher urease activity compared to other lineages, including CG258 and CG152 (TOP52). Interestingly, although we observed an overall increase in urease activity amongst our CG307 isolates, this did not correlate with enhanced fitness in artificial urine. Future studies will be critical for dissecting the role of *Kp* urease, and specifically the high levels of urease activity within CG307 strains, to UTI in a preclinical mouse model. Additionally, studies investigating the role of the high affinity UTS in the CG307 lineage in urea transport and breakdown will provide critical insights into whether urea metabolism contributes to establishing and/or maintaining UTI.

The strong association of CG307s with the urinary tract and the acquisition of several virulence factors likely to play a role in UTI suggested that the lineage may be adapted to this environment. Using a preclinical mouse UTI model, our results demonstrate that a representative CG307 strain – C234 – robustly colonizes the bladder. Strikingly, C234 bladder colonization phenocopied the UPEC strain UTI89, which is a well characterized cystitis isolate that results in robust UTI in the mouse model. Furthermore, C234 also colonized the bladder significantly better than TOP52. TOP52 has been extensively characterized for its *fimH*-dependent colonization of the bladder and ability to cause ascending infection of the kidneys in a mouse model of UTI [29,55]. Impactfully, the combined results of our study demonstrate that the CG307 strains outperformed TOP52 in nearly every assay, growing more effectively in AUM, exhibiting higher urease activity, and displaying significantly higher levels of bladder colonization. These observations suggest that while TOP52 caused symptomatic infection, the strain may not fully represent the lifestyle and pathogenic potential of CR*Kp* UTIs. Future work that seeks to determine the role the novel mechanisms that CG307 strains use to establish infection will provide critical insights into the pathogenic lifecycle of the epidemic CR*Kp* lineage currently circulating within the US.

In summary, this study reports that *Kp* UTIs are increasing in prevalence relative to UPEC across the Gulf Coast region. Additionally, the global epidemic lineage – CG307 – has continued to spread within the US and is now detected across the Gulf Coast. Characterization of CG307 strains indicate the lineage displays meaningful phenotypic adaptations that enhance growth in the urinary tract environment, increase urease activity, and enable robust UTI, which together may drive the observed increase in CG307 incidence across the Gulf Coast. CG307 strains decisively outcompete the established *Kp* UTI reference strain TOP52 (CG152), growing significantly better in AUM, exhibiting significantly higher urease activity, and colonizes the bladder at a much higher burden than TOP52. Impactfully, 1/3 of our CG307 isolates had a complete deletion of the *fim* operon. Yet, all of these strains were isolated from urinary tract, and in some patients, caused a symptomatic UTI indicating the CG307 lineage encodes a unique and yet unidentified means of establishing and maintaining UTI. These phenotypic adaptations are also likely to be driven by acquired factors including a high-affinity urea ABC transporter (UTS) and/or a glycogen synthesis operon (*glgBXCAP*). Future studies dissecting the mechanisms driving the rise in *Kp* UTI, and specifically MDR *Kp*, will provide critical insights into how *Kp* colonizes the urinary tract and results in increased morbidity, particularly in comparison to other non-MDR *Kp* isolates.

## MATERIALS AND METHODS

### Human Subjects Approval

Clinical isolates used in this study were collected previously [9] in accordance with the approval of the University of Texas Health Science Center at Houston Institutional Review Board, and the Committee for Protection of Human Subjects (CPHS; protocol ID HSC-MS-16-0334; ethical approval 5/16/2016), no further data were collected from these subjects. All isolates were collected after informed consent was obtained and were deidentified prior to use in this study. CG307 strain SL136 was collected by Dr. Allyson Shea’s group from excess clinical specimens acquired as part of standard clinical practice and associated microbiological and clinical data were obtained through a retrospective chart review approved by the University of South Alabama IRB (protocol #2178590) with a waiver of HIPPA authorization. Data from clinical databases were deidentified, date-blinded and collected solely for public health surveillance purposes.

### Clinical Database Analysis to Determine UTI Incidence

The two reference databases - HealthConnect Texas and USA Health - represent the Gulf coast region within the Greater Houston area and in Southern Alabama with partial overlap into Eastern Mississippi and Western Florida, respectively. Data were collected from 2020 to 2024 from HealthConnect Texas and 2017 to 2024 from USA Health. Data from HealthConnect was obtained using structured diagnosis codes (ICD-10 and SNOMED), laboratory results (LOINC, microbiological culture data), and encounter-level information, and did not include access to identifiable patient-level information. Our analysis used data from patients with documented UTI diagnoses with clinical or microbiological evidence that the cause of UTI was *Kp*. Data from USA Health were obtained using PathNet Microbiology: Statistical Reports within Cerner for the USA Health System. We extracted orderable procedure data (C Urine) and associated laboratory results, including organism identification fields (Report Organism/Response; Klepne or EC). The dataset was restricted to documented urine culture-positive cases with organism identification confirmed by MALDI-TOF. No identifiable patient-level information was included. To determine the prevalence of *Kp* UTIs across the Gulf Coast region, the percentage of *Kp* UTIs was normalized and compared against the incidence of UPEC UTIs reported between 2020 and 2024 (HealthConnect Texas) and between 2017 and 2024 (USA Health). Incidence over time was estimated by aggregating the total number of UPEC UTIs and *Kp* UTIs, then charting the % of *Kp* UTIs against the total number of UPEC UTIs. Significance over time was determined using a Chi-square test.

### Whole Genome Sequencing of Clinical Isolates

All isolates, except SL136, were previously sequenced using Illumina (short-read) and Oxford Nanopore (long-read) platforms and assembled [9]. SL136 was short-read sequenced as previously described [9]. Briefly, SL136 was grown in Luria Bertani (LB) broth (Fisher; catalog #DF0446-17-3) for 2-4 h with shaking at 37°C, then pelleted, and genomic DNA extracted using the Qiagen QIAamp DNA minikit (Qiagen; catalogue #51306) following the manufacturer’s protocol. Genomic DNA then underwent library preparation for short-read sequencing using the Illumina NextSeq 2000 sequencer. The SL136 genome was assembled as previously described [2]. Briefly, read quality assessment was conducted with Raspberry v0.3, trimming was conducted with Trimmomatic v0.39, SPAdes v3.15.2 was used for assembly, and assembly quality was assessed using Quast v5.2.0 [2]. Genome completeness was assessed with BUSCO v5.7.0 and CheckM v1.1.2 [56,57].

### Phylogenetic Analyses of Representative CG307 Strains

A reference-based phylogenetic analysis was performed using single-nucleotide polymorphisms (SNPs) identified from whole-genome sequencing data of our collected CG307 strains. Reads from each isolate were mapped to the internal C246 reference genome using Snippy [58] (v4.6), to perform read alignment and variant calling. To control for homologous recombination, the core SNP alignment was processed with Gubbins [59] (v3), which identified and masked recombinant regions. The resulting recombination-masked SNP alignment was then used to generate a maximum-likelihood phylogenetic tree with bootstrap values using RAxML [60] (v8.2.X), and the final tree was visualized and annotated using iTOL [61] (v7). We screened our whole genome sequencing dataset using BLAST for the presence of the following virulence factors: the UTS (UrtABC/AmiF operon), the *cp2* gene cluster, and the *glgX/glgB* glycogen synthesis genes. The ABRicate-v0.9.8 tool (https://github.com/tseemann/abricate) was used in parallel with Kleborate to query the Comprehensive Antibiotic Resistance Database (CARD) [62] to characterize antibiotic resistance determinants including KPC-2 and *bla*_CTX-M-15_ (accessed 2020-06-17). K and O locus typing was conducted using Kleborate’s KAPTIVE tool [63]. Copy number was inferred from the assemblies by counting the number of unique insertion sites across the chromosome and plasmids for each isolate. As we do not have long-read sequencing data for SL136, the copy number of *bla*_CTX-M-15_ was not assessed.

### Bacterial Strains and Growth Conditions

Isolates included in this study are listed in **Table 1**. LB broth and agar (Fisher; catalog #DF0445-17-4) was used to grow and maintain all strains. Cultures were grown overnight at 37°C and with shaking for liquid cultures with the exception of liquid cultures used for mouse infection, which were grown statically.

### Growth Curves

Overnight cultures were diluted 1:1,000 in fresh LB or artificial urine (recipe as previously described [26]) in a 96-well microplate (Cellstar; catalog #650185) in technical triplicate. The optical density at 600nm was measured every 15 minutes over a 24-hour time course in a Multi-Detection Microplate Reader (BioTek Synergy H2, WA, US) at 37°C with intermittent shaking. Data are reported as the average of two biological replicates with three technical replicates, error bars indicate the standard deviation.

### Mucoviscosity Assay

Mucoviscosity was measured as previously described [36]. Briefly, overnight cultures were normalized to OD_600_ = 1.0 using a Multi-Detection Microplate Reader (BioTek Synergy H2). The normalized culture was then pelleted by centrifugation at 1,000x*g* for 5 minutes and the OD_600_ value of the supernatant was measured again. The mucoviscosity value was then calculated as the supernatant OD / the normalized OD. Data is reported as the average of two biological replicates with three technical replicates, error bars indicate the standard deviation.

### Urease Activity Assay

Urease activity was measured as previously described [37]. Briefly, overnight cultures were spun for 5 minutes at 10,000 rpm, washed in 1X volume of PBS, and resuspended in urease broth, either with or without urea (Fisher; catalog #BP169-500). Next, 200 µL of each strain was transferred to a U-bottom 96-well plate (Greiner; catalog #650185), in triplicate, with a minimum of two biological replicates. The OD was measured at 415nm and 560nm to measure color change, which correlates with pH changes resulting from urease breakdown of urea, and at 600nm to measure growth. ODs were measured every 20 minutes for 18 hours while shaking at 37°C in a microtiter plate reader (BioTek Synergy H1). The color change of the urea positive cultures was then normalized to the urea negative controls to account for any color change (pH change) that was not due to urease activity. Bar chart data is reported as the average of three technical replicates and includes two biological replicates, error bars indicated the standard deviation.

### Mouse Model of MDR *Kp* UTI

The mouse UTI model was performed as previously described [29]. Briefly, C57BL/6 female mice (Charles River Labs) were transurethrally infected with ∼2 x 10^7^ CFUs of bacteria. Bacterial inoculums for both *Kp* and UPEC were prepared by subculturing overnight cultures grown statically for 24 hrs at 37°C 1:250 in fresh LB. Subcultures were then grown statically at 37°C for an additional 24 hrs. Inoculums were then diluted 1 to 1 in 1X PBS, then centrifuged at 10,000 rpm for 10 minutes, washed with 1X PBS, resuspended in 1X PBS, and diluted to the desired CFU of 2×10^7^. Organs, including the bladder and kidneys, were harvested 1 day post infection (dpi) and homogenized using an MP-BIO Fastprep24 bead beater. Bacterial loads were enumerated following serial dilution onto LB plates and incubation at 37°C overnight. All animal work was approved by the Institutional Animal Care and Use Committee (IACUC) and Animal Welfare Committee (AWC) at the University of Texas Health Science Center at Houston (protocol # AWC-23-0049). Data includes two biological replicates with five mice per replicate.

### Statistical Analyses

All *in vitro* experiments included at least two biological and three technical replicates. To determine significant differences in growth between strains, the area under each curve was determined, and normality was tested with a Shapiro-Wilk test. Statistical comparisons between groups were performed using the Kruskal-Wallis test, with Dunn’s post-hoc test used for pairwise comparisons between groups. We then used a Kruskall-Wallis test to determine variance between groups and determined variance between groups using a Dunn’s post-hoc pairwise test. To determine significant differences in urease activity between strains, urease activity was graphed over time for 18 hours, area under each curve analyses were determined, and a Mann-Whitney U Test was used to compare activity between strains. Our *a priori* Wilcoxon rank-sum sample size calculations determined a minimum of 8 mice per experimental group were required using a beta value of 0.8 using an alpha value of 0.05. Each mouse is graphed individually, and the median of each group is denoted by the black bar. The Wilcoxon rank-sum test was also used to determine significant differences between strains. To test for the effects of clonal group or site of isolation in our mucoviscosity and urease activity data respectively, strains were first grouped, then compared using a Linear Mixed Effects Model (LMM) if comparing three or more groups, or with a Chi-squared test if comparing only two groups. To evaluate whether the relative frequency of *Kp* UTIs increased over time compared to *E. coli*, a generalized linear model with a binomial distribution (logistic regression) was fit, modeling organism identity (*Kp* vs. UPEC) as a binary outcome and calendar year as a continuous predictor. RStudio v2025.09.1+401 and GraphPad Prism v10.6.1 were used for figure creation and statistical analysis; *p =* *: <0.05.

## ACKNOWLEDGEMENTS

We thank Dr. Scott Hultgren for the TOP52 and UTI89 strains [64]. This work was supported by an NIH K01 DK128381-01A1 (JNW), the University of Texas STAR award (JNW), an NIH P01 AI152999-01 (BMH), the University of South Alabama College of Medicine Start-up funds (AES), a Fullbright Visiting Scholar Award (EV), and an American Heart Association Predoctoral Fellowship (JMDR).

## AUTHOR CONTRIBUTIONS

KDB: project development, data collection and analysis, manuscript writing and editing. JMDR: project development, data collection and analysis, manuscript writing and editing. JG: data collection and analysis, manuscript writing and editing. TRC: data collection and analysis. EV: data collection and analysis, manuscript writing. MNS: data collection and analysis. AES: data collection and manuscript writing. JNW: data collection and analysis, project development, and manuscript writing and editing. BMH: data collection and analysis, project development, manuscript writing and editing. All authors contributed to the article and approved the submitted version.

## DATA AVAILABILITY

All short-read and long-read FASTQ files along with complete assemblies for all *K. pneumoniae* isolates sequenced, except SL136, are publicly available and can be found in BioProject PRJNA658369. SL136 fastq and fasta files can be found in BioProject PRJNA1398917, BioSample SAMN54466988. Relevant R code is available at https://github.com/Hanson-Lab/publications/Kp_CG307.

## ETHICS DECLARATIONS

All bacterial isolates were collected after informed consent was obtained and were deidentified prior to use in this study. The collection of bacterial isolates was approved by the University of Texas Health Science Center at Houston Institutional Review Board, and the Committee for Protection of Human Subjects under protocol HSC-MS-16-0334. Strain SL136 was isolated from excess clinical specimens collected as part of routine care. A retrospective chart review was conducted under University of South Alabama IRB protocol #2178590 with a waiver of HIPAA authorization, prior to being provided to study investigators. Additionally, collection of isolates was performed in accordance with the 1964 Helsinki declaration and its later amendments.

## APPROVAL FOR ANIMAL EXPERIMENTS

All animal work was approved by the Institutional Animal Care and Use Committee (IACUC) and Animal Welfare Committee (AWC) at the University of Texas Health Science Center at Houston (protocol # AWC-23-0049).

